# When morphology stands still: constrained floral evolution in a mega-diverse legume genus

**DOI:** 10.1101/2025.11.07.687212

**Authors:** Monique Maianne, Yago Barros-Souza, Fabio A. Machado, Leonardo M. Borges

## Abstract

- Changes in floral morphology are key for the evolutionary success of flowering plants. However, many species-rich and ecologically diverse groups show low variation in floral traits. Low morphological variation may be associated with selective pressures and constraints to changes. Nonetheless, because our understanding of floral morphological evolution is focused on plant groups with diverse and specialized floral morphology, there has been little progress in identifying factors underlying floral morphological uniformity.
- Herein we addressed this issue by asking whether the morphology of flowers in the mega-diverse legume genus *Mimosa* is shaped by constraints. By investigating patterns and processes underlying trait variation along floral whorls, we evaluated the role of development and/or function in limiting morphological change.
- Our results showed that floral morphology is remarkably homogeneous and likely shaped by convergent evolution. Morphological variation is unevenly distributed across floral whorls, reflecting differences in the levels of constraints.
- Floral morphology in *Mimosa* is unlikely to have been primarily driven by pollinator-mediated selection. Instead, we suggest that the development and function of flowers in the mimosoid inflorescence constrained morphological evolution in this megadiverse yet morphologically uniform group.

## INTRODUCTION

Flowers are astonishingly diverse in size, shape, color, scent and reproductive strategies (Endress, 2011; Harder and Barrett, 2006). Like other reproductive structures (e.g., Sauer and Hausdorf, 2009; Simmons and Firman, 2014; Stefanini et al., 2021; Thomaz et al., 2019), flower morphology is subject to strong selective pressures due to its impact on offspring production (Harder and Johnson, 2009; Vamosi et al., 2018). Even small changes in genetics (Lüthi et al., 2022), development (Endress, 2011), pollinator community (Cardona et al., 2020), or other ecological factors (e.g., floral antagonists (Huang et al., 2022)) can lead to changes in floral morphology and influence diversification rates (Huang et al., 2022; Ronse De Craene, 2018; Vamosi et al., 2018). Nevertheless, the ecological (widespread distribution) and evolutionary (high species diversity) success of groups with superficial uniform floral morphology (e.g., Barneby, 1991; Chartier et al., 2021; Vasconcelos et al., 2019) suggests that restrictions on morphological evolution may be stronger than the pressures to change (Smith et al., 1985; but see Davis et al., 2014). Whether these limitations play a major role in shaping floral morphology and to what extent different floral traits are affected by them in such highly diverse groups of plants remains unclear.

Flowers are complex phenotypes composed by different organs (sepals, petals, stamens, and carpels) that develop through distinct ontogenetic pathways and may fulfill different functions in plant reproduction (Endress, 2011; Shan et al., 2019). As such, each organ (or set of organs) may be subjected to distinct evolutionary constraints (Diggle, 2014; Walker, 2010), which could lead to differences in the degree of morphological variation among floral whorls (Chartier et al., 2017; Ordano et al., 2008; Rosas-Guerrero et al., 2011). Investigating how morphological variation – i.e., disparity – deviates among whorls and how it has been shaped over time may explain how constraints shape the evolution of flowers. Answering this question is particularly relevant to understand why and how flowers of so many groups remain morphologically stable (e.g., American Malpighiaceae (Davis et al., 2014); Brassicaceae (Gómez et al., 2016); Leandra Raddi (Melastomataceae) (Reginato and Michelangeli, 2016); Myrcia DC. ex Guill. (Myrtaceae) (Vasconcelos et al., 2019); Sapotaceae (Chartier et al., 2017) and *Mimosa* L. (Leguminosae, Caesalpinioideae) (Barneby, 1991)).

Despite being the fifth largest genus of legumes and one of the most diverse genera in the Americas (Barneby, 1991; Lewis et al., 2005; Simon et al., 2011), *Mimosa* flowers seem to have remained relatively similar throughout ≈ 24 million years of evolution. This is particularly surprising because *Mimosa* lineages occur in areas highly diverse in pollinators, which could pro-mote floral disparification (e.g., Aguiar et al., 2024; Nava-Bolaños et al., 2023). Yet, there is no clear floral morphological specialization among *Mimosa* lineages, even between species that are pollinated by widely different functional groups, such as bats and bees (Barneby, 1991; Silva et al., 2011; Vogel et al., 2005). Flowers in the 600+ *Mimosa* species are radially symmetric, have a tubular and reduced perianth, exposed and showy filaments and styles, and, like other mimosoid legumes, are densely aggregated in capitate or spicate inflorescences (Barneby, 1991; Borges et al., 2024; Koenen et al., 2020; Tucker, 2003).

Because *Mimosa* floral bauplan is shared over space and time across *Mimosa* lineages, we hypothesize that flower evolution is constrained, probably by development. Whereas non-mimosoid legumes frequently show different patterns of organ initiation per whorl, dissimilar differentiation in organs of the same whorl, and variations in the temporal overlap of initiation across whorls (Tucker, 2003), these features are comparatively more stable in mimosoids. Flowers in *Mimosa* and in the majority of mimosoids develop synchronously along the inflorescence and, within flowers, organs of the same whorl are alike and initiate simultaneously (except sepals) (Gonçalves et al., 2024; Pedersoli et al., 2023; Tucker, 2003). Because changes in the timing and rate of developmental events are among the main drivers of floral morphological diversity (Naghiloo, 2020; Ronse De Craene, 2018), the absence of such variations may hinder morphological diversification. However, phylogenetic comparative studies investigating floral evolution in mimosoid flowers are still pending.

Herein, we quantified the morphological variation and investigated the evolution of flowers across phylogenetically and geographically representative species of the mega-diverse genus *Mimosa*. Using Phylogenetic Comparative Methods (PCMs), we integrated morphological data with a comprehensive phylogeny to investigate the underlying evolutionary processes shaping floral disparity to address three questions: (1) Is floral morphology of different *Mimosa* lineages more similar than expected given their phylogenetic relatedness, indicating that floral morphological diversity has been shaped by evolutionary convergences?; (2) Do disparity varies among floral whorls, as expected if they are subject to different evolutionary constraints?; (3) Is the evolution of floral whorls biased toward a morphological optimum, as expected if constraints hinder morphological change?

## MATERIAL AND METHODS

### Sampling and morphometrics

We sampled 15% of the taxonomic diversity of *Mimosa* (83 species). Sampling was defined to capture the broadest range of phylogenetic, geographic and floral diversity of the genus (Barneby, 1991; Borges et al., 2024; Simon et al., 2011). *Mimosa*’s phylogeny shows high geographical structure and includes 24 main clades with strong phylogenetic support (Simon et al., 2011). At least 10% of the diversity of each of these clades was sampled here.

For each species, we collected 14 quantitative floral traits from herbarium specimens (Figure 1; see Table S1 for a full list of vouchers). We focused on morphological floral traits that describe each floral whorl, vary among *Mimosa* flowers, and could be consistently recorded across all species. For each species, we selected approximately three flowers from the apex of an inflorescence. The flowers were rehydrated, and each floral organ was mounted in glycerin jelly on a glass slide. We then photographed each floral organ using digital cameras mounted on stere-omicroscopes. To measure flowers, we placed landmarks describing each floral traits using the TPS Dig program (Rohlf, 2006) and then obtained linear measurements based on the landmarks with the *geomorph* package (Adams and Otárola-Castillo, 2013). Finally, we averaged each floral trait across flowers within the same inflorescence, and also across individuals in species for which we sampled more than one voucher from different localities. All variables were logtransformed before analyses.

**Figure 1:**
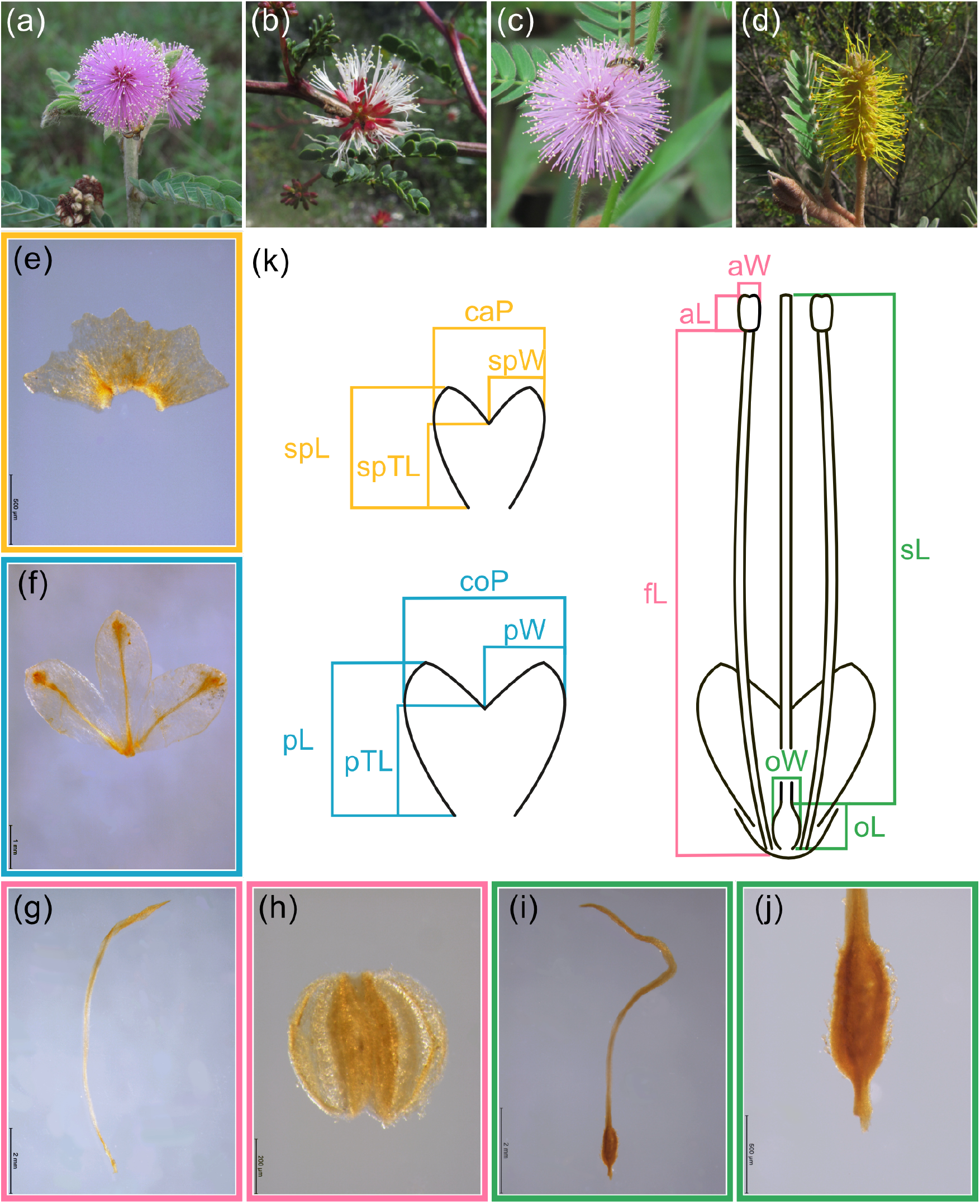
Schematic and photographic representation of *Mimosa* floral morphological diversity. (a)-(d): Photographs of inflorescences of *M. bombycina* Barneby, *M. pringlei* S.Watson, *M. hirsutissima* Mart. *M. barretoi* Hoehne (from left to right). (e)-(j): Photographs of floral structures of *M. minarum*, exemplifying the images used to extract linear measurements. (e): calyx; (f): corolla; (g): filament; (h): anther; (i): carpel; (j): ovarium. (k): Schematic illustration of the measured floral traits. caP: calyx perimeter; spW: sepal width; spTL: sepal tube length; spL: sepal length; coP: corolla perimeter; pW: petal width; pTL: petal tube length; pL: petal length; fL: filament length; aW: anther width; aL: anther length; sL: style length; oW: ovary width; oL: ovary length; Yellow indicates calyx traits; blue, corolla traits; pink, androecium traits; green, gynoecium traits.

### Floral phylomorphospace and disparity

To visualize the floral morphospace, we performed a Principal Component Analysis (PCA) on the covariance matrix (*PCA* function; *FactoMineR* package Lê et al., 2008) and projected the *Mimosa* phylogeny (Vasconcelos et al., 2020) onto this morphospace (*phylomorphospace* function; *phytools* package; Revell, 2012).

To investigate how floral disparity is distributed among floral whorls in *Mimosa*, we first followed Chartier et al. (2017). We estimated Euclidean distance matrices for all floral traits in the dataset, as well as separately for each floral whorl (*dist* function; *stats* package). To investigate which floral whorl contributes the most to total floral disparity, we performed Mantel tests between the Euclidean distance matrix for the entire floral dataset and those for each individual floral whorl (*mantel* function; *vegan* package). To determine whether the disparity of individual floral whorls are greater or less than the disparity of the total floral dataset, we compared the mean pairwise Euclidean distance between taxa of each floral whorl with the mean pairwise Euclidean distance of 1,000 randomly resampled morphological matrices. The randomizations were done considering the same number of traits as in the subset being compared. A whorl was considered more or less variable than the rest of the flower if its mean pairwise distance was higher or lower than 97.5% of those of the resampled matrices.

We also estimated the variance-covariance matrix to access the amount of variation of each floral trait within a whorl. To estimate the confidence intervals for the variances of floral traits, we obtained 1,000 matrices from a posterior distribution generated using a Bayesian model for matrix estimation using a conjugate inverse Wishart prior (*BayesianCalculateMatrix* function; *evolqg* package; Melo et al., 2015).

### Phylogenetic signal and evolutionary models

To investigate evolutionary patterns associated with *Mimosa* floral disparity, we first tested whether floral trait values are distributed across species according to their phylogenetic relatedness. To do so, we measured the phylogenetic signal of each floral whorl using a multivariate version of the Blomberg’s *K-statistic* (Adams, 2014; Blomberg et al., 2003) (*physignal* function; *geomorph* package; Adams and Otárola-Castillo, 2013). If *K* equals 1, then the distribution of trait values across the phylogeny is influenced by phylogenetic relatedness as expected under a Brownian Motion model (BM). *K* values lower than 1 suggest that distantly related species are more similar than expected under BM, whereas *K* values greater than 1 indicate that closely related species are more similar than expected under a BM (Adams, 2014; Blomberg et al., 2003). We measured the statistical significance of *K* randomizing trait values among species 999 times.

Phylogenetic signal can inform about the consequences of evolutionary processes but not about the processes of morphological evolution themselves (Revell et al., 2008). To model morphological evolution of each floral whorl, we fitted two multivariate models, the neutral model, BM (Felsenstein, 1985), and the adaptive model, Ornstein-Uhlenbeck (OU) (Hansen, 1997, 2014) (mvBM and mvOU function; *mvMORPH* package; Clavel et al., 2015). Then, to understand the role of evolutionary constraints on floral whorls and their traits, we focused on transformations of OU parameters. Specifically, we estimated the phylogenetic half-lives (*t*1*/*2) and the stationary variances (*vy*) (Butler and King, 2004; Hansen, 1997; Hansen et al., 2008). *t*1*/*2 were calculated as ln(2)*/α* (Hansen et al., 2008), being the *α* the diagonals of the selective rate matrices *A* (Bartoszek et al., 2012). *t*1*/*2 estimate the time needed for a lineage to reach halfway between the ancestral state and the regime optimum (Hansen et al., 2008). Higher *t*1*/*2 suggests relaxed attraction of the ancestral state toward the regime optimum, whereas lower *t*1*/*2 indicates a strong attraction toward the optimum and, thus, greater evolutionary constraints (Grabowski et al., 2023; Hansen, 2014; Hansen et al., 2008). *vy* matrices was calculated using the stochastic rate matrices Σ and the *A* matrices following Bartoszek et al., 2012. Then, we accessed the value of *vy* of each trait within an whorl from the diagonals of the *vy* matrices. High *vy* suggest a large amount of residual variance not explained by the strength of attraction to the primary optima, whereas low *vy* indicates that the variance is well represented by the pull of attraction toward the primary optima (Hansen, 1997).

We measured the uncertainty around evolutionary model parameters by sampling points around their maximum likelihood using the *dent_walk* function; *dentist* package (Boyko and O’Meara, 2024). The best evolutionary models explaining traits distribution of floral whorls were selected based on the Akaike Information Criterion corrected for small sample sizes (AICc).

## RESULTS

### Floral phylomorphospace and disparity

The first two axes of the phylomorphospace explained 73.97% of the total floral morphological variance (Figure 2A; Table S2). The first Principal Component (PC) accounted for 54.24% of morphological variation (Figure 2A), which was mainly attributed to long and wide versus short and narrow calyxes and contrasting lengths of corolla tube (Table S2). The second PC accounted for 19.73% of the morphological variation (Figure 2A) and was characterized by long versus short styles, filaments and sepals (Table S2). The projection of the phylogeny into the PC graphs, i.e., the phylomorphospace, showed a strong overlap among distantly related lineages and long branches between them (Figure 2A). Although variation within the calyx was more spread along PC axes than in other whorls, the overlap of lineages and lack of a clear structure in the phylomorphospaces were also observed for each floral whorl (Figure 2B-E). Taken together, these patterns of phylomorphospace occupation for the whole flower and individual whorls are indicative of morphological convergences (Figure 2B-E); see also Figure S1 for representations in other PC axes).

**Figure 2:**
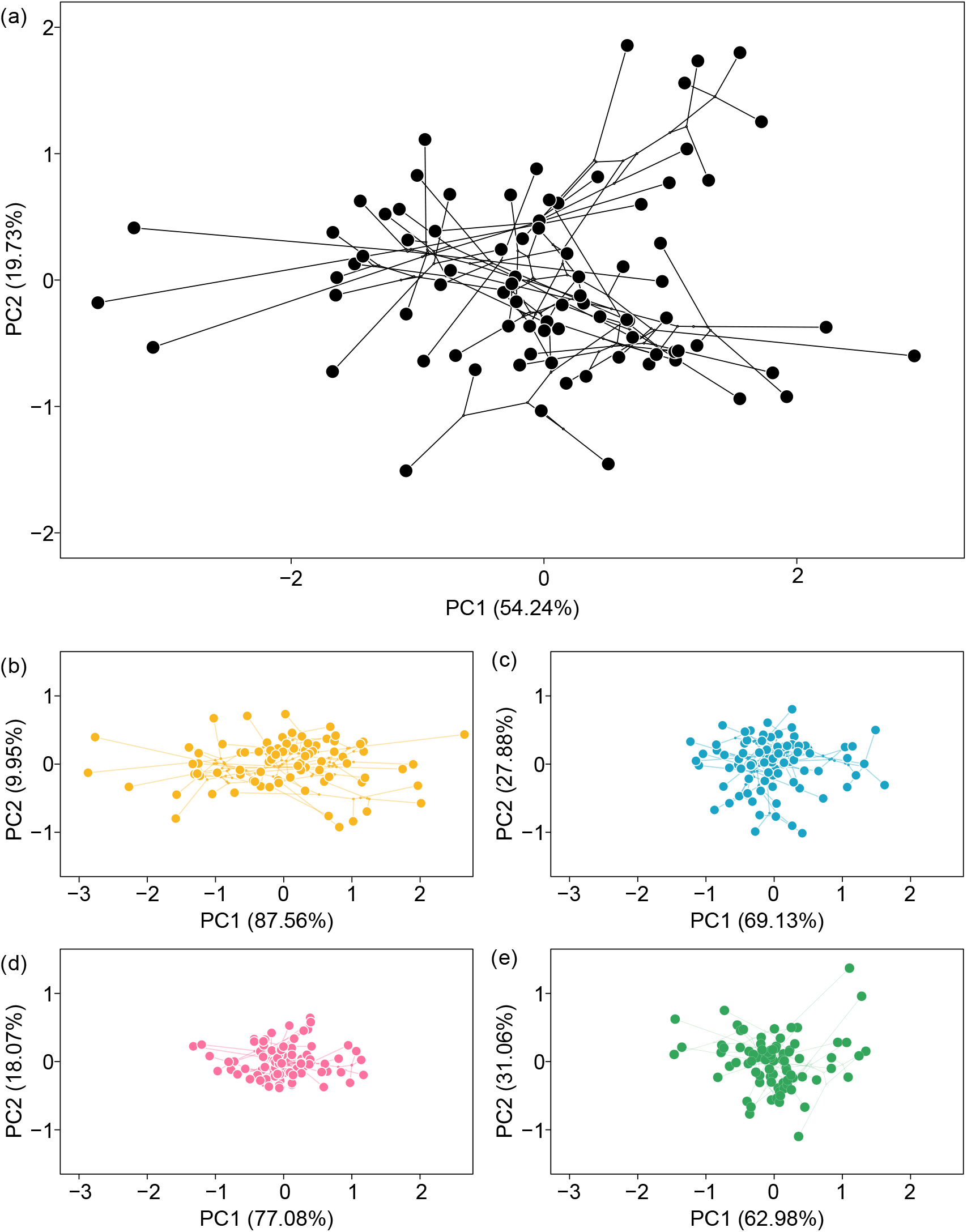
Phylomorphospaces based on the first two Principal Components (PC) of floral morphological diversity in *Mimosa*. (a) Phylomorphospace depicting all floral morphological traits; (b) calyx phylomorphospace; (c) corolla phylomorphospace; (d) androecium phylomorphospace; and (e) gynoecium phylomorphospace.

Consistent with the variance contributions within the PCs, the disparity analyses showed high morphological variation in the calyx. It was the whorl most correlated with total floral disparity (*r* = 0.79, *p <* 0.005) and exhibited the highest mean pairwise distance between species (*MPD* = 1.3), followed by the corolla (*r* = 0.69, *p <* 0.005, *MPD* = 0.89), gynoecium (*r* = 0.59, *p <* 0.005, *MPD* = 0.84), and androecium (*r* = 0.63, *p <* 0.005, *MPD* = 0.69). The observed calyx disparity was greater than 95.7% of resamplings of other floral traits, while the observed disparity of the corolla and androecium were lower than the values of 82.2% and 91.7% of the null distributions, respectively (Figure 3). The gynoecium showed no significant differences compared to the rest of the flower (Figure 3).

**Figure 3:**
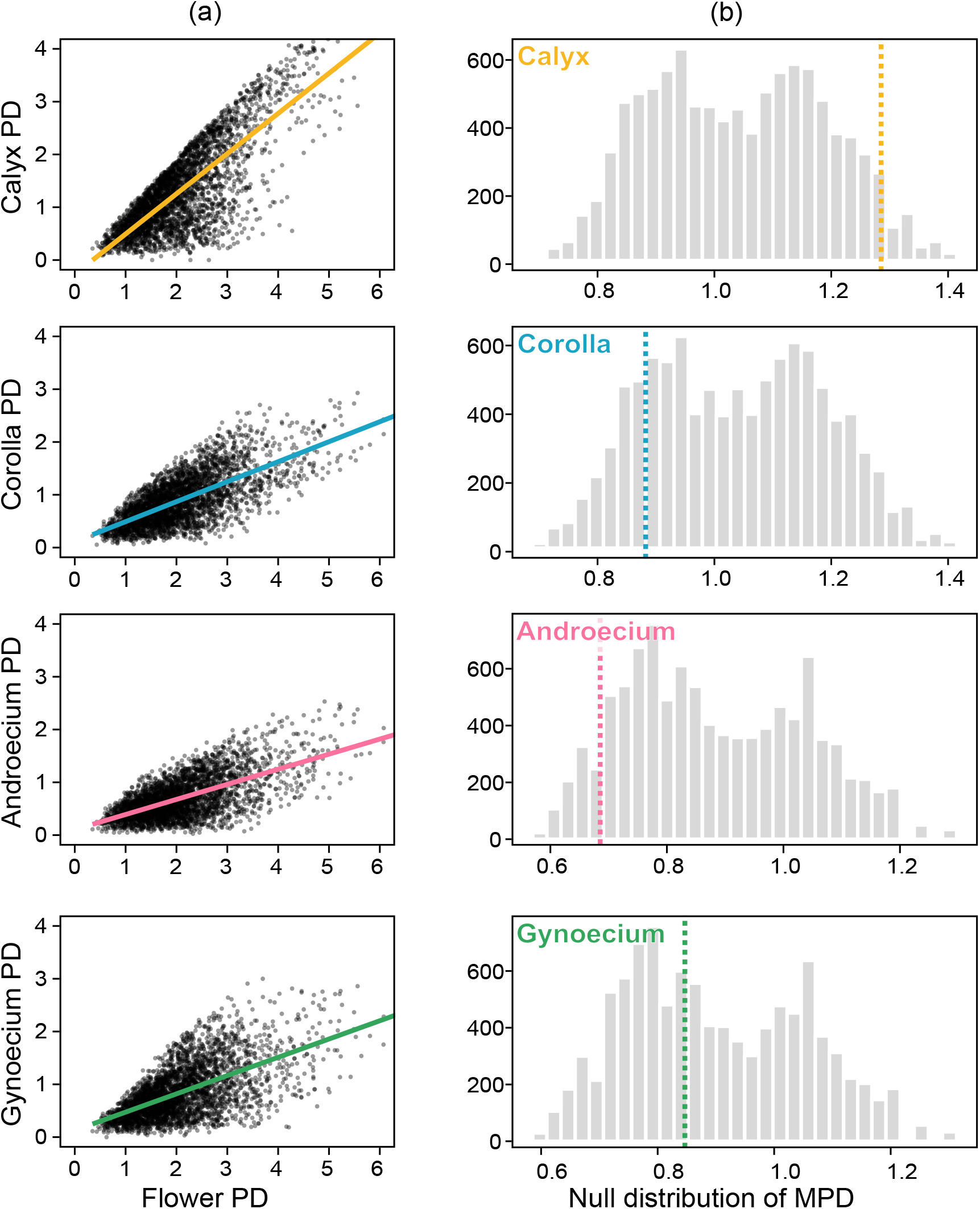
Floral morphological variation based on Euclidean pairwise distances (PD) between *Mimosa* species. (a) Correlation between the pairwise distances of each floral whorl (y-axis) and the pairwise distances of the total floral dataset (x-axis). Lines indicate the linear regression between the two sets of PDs. (b) Null distributions of the mean pairwise distances (MPD) for each floral whorl. Dashed lines represent the observed mean pairwise distances estimated for each floral whorls. In yellow, the results for the calyx, in blue for the corolla in pink for the androecium, and in green for the gynoecium.

The variance-covariance matrix showed the the same trend of disparity among floral whorls reported just above. The calyx has the greatest variance (*Var* = 1.19), followed by the corolla (*Var* = 0.51), the gynoecium (*Var* = 0.48), and the androecium (*Var* = 0.32). The amount of variance in each whorl was distributed unevenly among traits (Figure 4). Within the calyx, the variance of the sepal length was greater than calyx perimeter and sepal width. In the corolla, petal tube length had the greater variance, and the length of filaments and styles showed the grater variance within the androecium and gynoecium, respectively (Figure 4). All these variances fall within the confidence intervals estimated for the sample from the posterior distributions (Figure 4).

**Figure 4:**
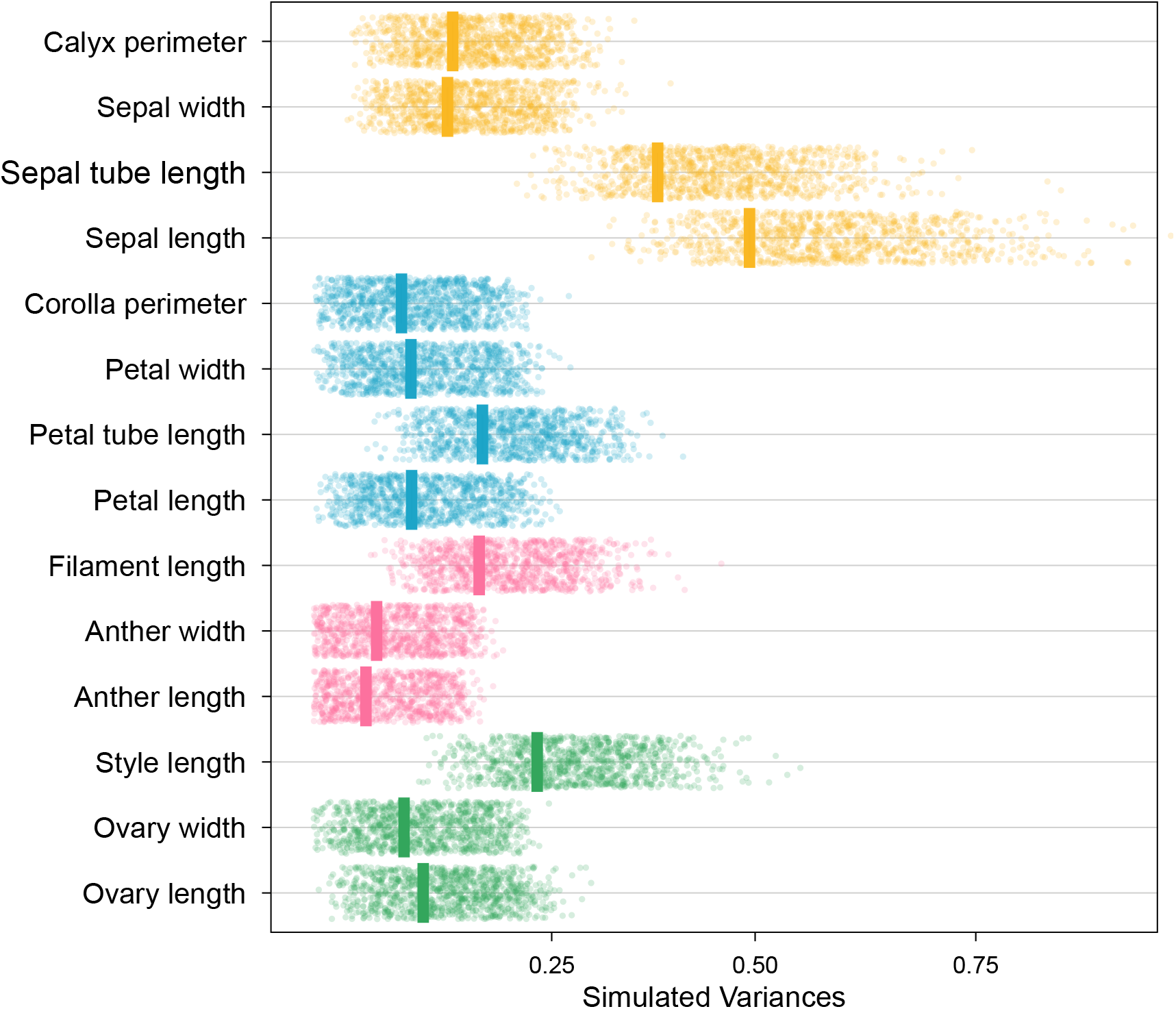
Variances of each floral traits. Dots represent the variance estimated from the posterior distributions and vertical bars indicate the observed variance of each floral trait. In yellow, calyx traits, in blue corolla traits, in pink androecium traits, and in green gynoecium traits.

### Phylogenetic signal and evolutionary models

Following the lack of phylogenetic structure in the phylomorphospaces, all floral whorls exhibited low phylogenetic signal, suggesting that the morphology of distantly related species is more similar than expected under a Brownian Motion (BM) model of evolution (*Kmult <* 1). Only the *Kmult* estimated for the gynoecium was not statistically significant at *p <* 0.05 (Table 1). Additionally, the adaptive Ornstein-Uhlenbeck (OU) models provided a better fit than the BM models for all floral whorls (Table 1), again, showing that the phylogenetic relatedness among species is not the primary factor structuring floral morphological variation. Instead, the floral morphology in *Mimosa* evolved under evolutionary pressures toward morphological optima (Table 1).

**Table 1:** Disparity indices and results of evolutionary models for each floral whorl in *Mimosa*. MPD: mean pairwise distance. K-mult: multivariate phylogenetic signal.

|  | **Calyx** | **Corolla** | **Androecium** | **Gynoecium** |
| --- | --- | --- | --- | --- |
| **Disparity** |  |  |  |  |
| MPD | 1.3 | 0.89 | 0.69 | 0.84 |
| Variance | 1.19 | 0.51 | 0.32 | 0.48 |
| **K-mult** |  |  |  |  |
|  | 0.44 | 0.43 | 0.42 | 0.22 |
| **Brownian-Motion** |  |  |  |  |
| AICc | 2.912 | -153.43 | 17.85 | 266.91 |
| loglik | 13.21 | 91.38 | 0.45 | -124.08 |
| **Ornstein-Uhlenbeck** |  |  |  |  |
| AICc | -53.84 | -205.54 | -25.51 | 160.34 |
| loglik | 52.87 | 128.74 | 28.79 | -64.16 |
| Phylogenetic half-life | 0.18 | 0.3 | 0.04 | 0.007 |
| Stationary variance | 1.28 | 0.86 | 0.29 | 0.46 |

Phylogenetic half-lives (*t*1*/*2) and stationary variances (*vy*) differed markedly among floral whorls and within each whorl (Figure 5, Table 1). The corolla showed the highest mean of phylogenetic half-lives, followed by the calyx, androecium and gynoecium (Table 1). Calyx summed the greatest stationary variance values, followed by the corolla, gynoecium, and the androecium (Table 1). Within the calyx, traits describing the vertical variation presented lowest *t*1*/*2 and highest *vy* (Figure 5). Although the corolla was less morphologically variable than the calyx, traits describing horizontal variation exhibited high *vy* values (Figure 5). These high *vy* values were associated with their highest *t*1*/*2, indicating that the weak attraction of these traits to a morphological optimum allowed greater accumulation of morphological variation. The length of the filaments and styles showed the highest *vy* values, but similar *t*1*/*2 compared to other traits within the androecium and gynoecium, respectively (Figure 5). Altogether, the low values of *t*1*/*2 and *vy* in the androecium, gynoecium and in traits describing the length of the corolla organs indicate less morphological variation associated with a strong pull of attraction of these traits toward morphological optima, while the other floral traits are less evolutionarily constrained.

**Figure 5:**
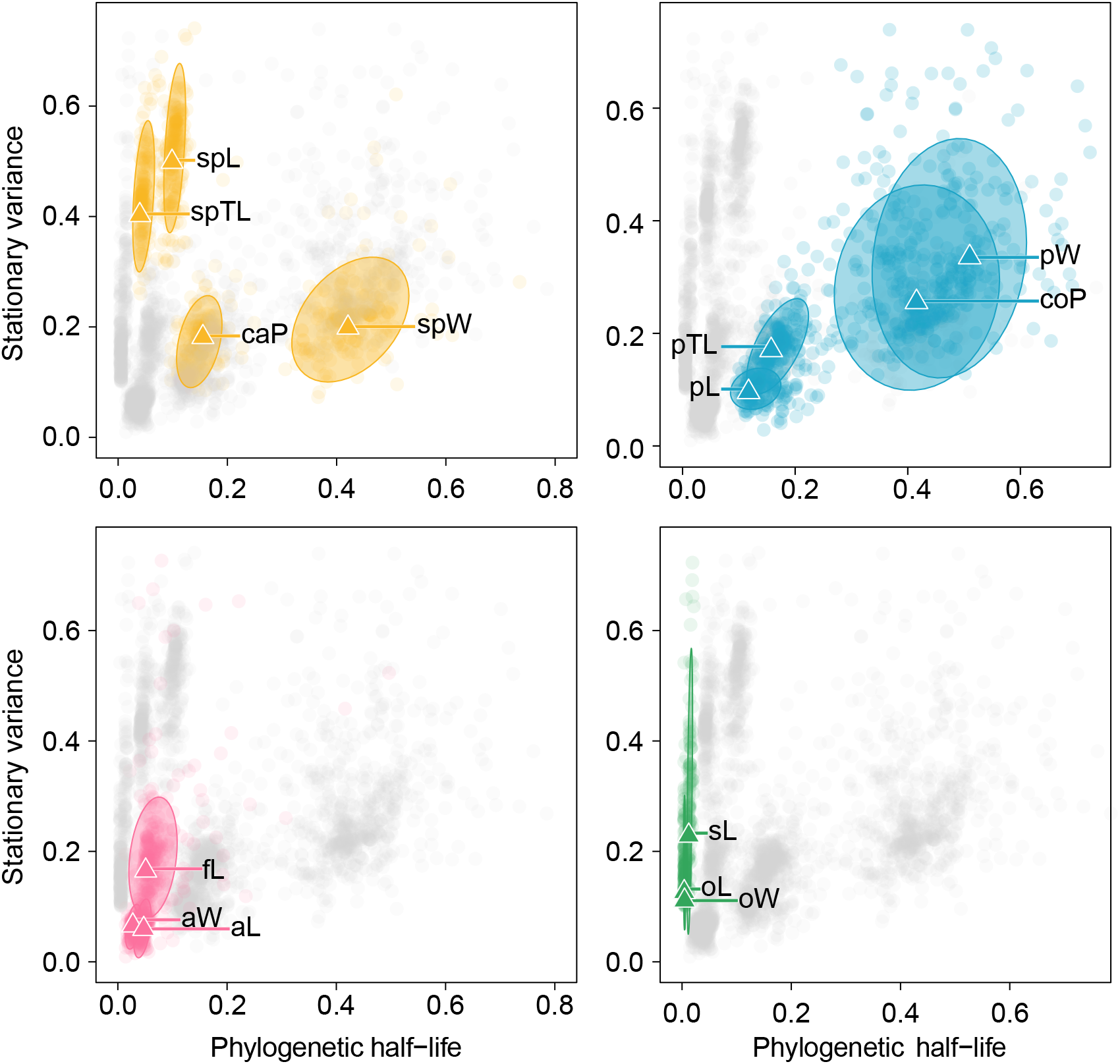
Parameters of evolutionary models estimated for each trait within *Mimosa* floral whorls. Triangles represent the observed values of stationary variance (y-axis) and of phylogenetic half-life (x-axis). Dots represents the sampling points around maximum likelihood estimates and ellipses around triangles represents the confidence interval (95%). In yellow, results for the calyx, in blue for the corolla, in pink for the androecium, and in green for the gynoecium. caP: calyx perimeter; spW: sepal width; spTL: sepal tube length; spL: sepal length; coP: corolla perimeter; pW: petal width; pTL: petal tube length; pL: petal length; fL: filament length; aW: anther width; aL: anther length; sL: style length; oW: ovary width; oL: ovary length.

## DISCUSSION

Given the influence of floral morphology on the reproductive success of angiosperms, understanding the origins of floral variation is crucial to pinpoint the drivers of diversity and evolutionary success in flowering plants (Chartier et al., 2014; Sauquet and Magallón, 2018). Here, we quantified the variation of floral morphology —floral disparity— through different approaches and investigated the evolutionary processes that shaped this disparity in *Mimosa*, a mega-diverse genus of legumes. Our results showed that floral morphology is remarkably homogeneous and likely shaped by convergent evolution. Below, we argue that this pattern is unlikely to have been primarily driven by pollinator-mediated selection. Instead, we suggest that the development of flowers in the mimosoid inflorescence constrained floral evolution in this group.

### On the link between pollination and convergence

All floral whorls showed low phylogenetic signal (Table 1), indicating that floral morphology of distantly related species are more similar to one another than expected by their evolutionary relatedness (Adams, 2014; Revell et al., 2008). The overall branch overlap and clustering of distantly related *Mimosa* lineages in the phylomorphospace strongly suggest a pattern of morphological convergence (Figure 2), defined by lineages evolving to be more similar to one another than their ancestors (Losos, 2011). Convergence is typically associated with shared selective pressures (Losos, 2011; Mayr, 1963; Simpson, 1953), which lead distantly related lineages to-wards similar adaptive peaks. Indeed, our results show that all floral whorls fit an Orstein-Uhlenbek (OU) model of evolution (Table 1), indicating that traits within each floral whorl are constrained to an adaptive optimum (Butler and King, 2004; Hansen, 1997, 2014). In the context of animalpollinated flowers, as those of *Mimosa*, limited disparity often arises from stabilizing selection driven by the interactions with pollinators (e.g., Artuso et al., 2021; Gómez et al., 2016; Joly et al., 2018; Lagomarsino et al., 2017; Simón-Porcar et al., 2024), where long-term floral morphological stasis is driven by the stability of the pollination niche (Davis et al., 2014; Vasconcelos et al., 2019).

Pollination niches are still largely unrecorded for most *Mimosa*. Nonetheless, existing evidence indicates that some species are pollinated by a single functional group (e.g., bats [Vogel et al., 2005]), while others are more generalist (e.g., visited by flies and small bees [Amorim et al., 2013; Borges et al., 2017; Silva et al., 2011]; butterflies, small bees, and bumblebees [our own observation]). These evidences and the high diversity of pollinators across the broad Neotropical distribution of *Mimosa*, suggest that a stable pollination system is unlikely to be the primary driver of floral convergence. Further evidence supporting this idea comes from functional morphology. For plants in which convergences are associated with pollinators, the main axes of morphological divergence comes from traits directly involved in pollinator interactions (e.g., Rose and Sytsma, 2021). However, for *Mimosa*, trait variation accumulates mainly in the calyx, a whorl not directly involved in pollination of these plants (Silva et al., 2011; Vogel et al., 2005). Pollination in *Mimosa* occurs mainly via interaction with the stamens (and styles), which are attractive and may serve as a landing platform for pollen harvest across the densely packed inflorescence (e.g., Figure 1c). In some cases, non-structured nectaries provide a resource for larger pollinators (e.g. bats and hummingbirds), which, again, are attracted by the display and maybe also scent of the less variable reproductive whorls (de Sousa et al., 2025; Vogel et al., 2005). If pollinator interactions are not the primary drivers of morphological evolution in *Mimosa* flowers, then other factors are needed to explain its disparity.

### Disparity of floral whorls aligns with development

The uneven distribution of extant morphological variation within the four floral whorls (Figure 3) aligns with floral developmental patterns in *Mimosa*.

Highest morphological variation in the calyx reflects its greater developmental variability. In *Mimosa*, and other mimosoid legumes, sepals vary in (1) order of initiation (parts of all other whorls initiate simultaneously); (2) elongation rate, as sepals may elongate unevenly; and (3) time of establishment of gamosepaly, with sepals uniting either early or late in development (Gonçalves et al., 2024; Prenner, 2004; Ramírez-Domenech and Tucker, 1990, 1989; Tucker, 1992). Together, these developmental features interact to create the strikingly different calyx morphologies seen in *Mimosa*, which vary from a simple shallow ring at the base of the corolla to a short cup with long, deeply fringed lobes (Barneby, 1991).

The comparatively low disparity of other whorls also reflects their developmental features. In the corolla, petals of *Mimosa* and other mimosoids initiate simultaneously as free and equidistant primordia, elongate uniformly, and enclose the androecium and gynoecium in valvate aestivation (Gonçalves et al., 2024; Pedersoli et al., 2023). These features make the corolla one of the most developmentally stable organs of mimosoid flowers (Gonçalves et al., 2024; Pedersoli et al., 2023; Prenner, 2004; Ramírez-Domenech and Tucker, 1990). This pattern differs from that of other caesalpinioid legumes in which greater developmental plasticity in the corolla is linked to increased floral disparity (e.g., Bruneau et al., 2014; Marazzi and Endress, 2008; Tucker, 1996; Zimmerman et al., 2013).

The next whorl to arise during development, the androecium, was the least disparate in *Mimosa* flowers. Again, this differs widely from other groups of legumes and angiosperms, in which the androecium is one of the most variable floral organs (Chartier et al., 2017; Herting et al., 2023; Sinjushin, 2021). In other angiosperms, greater androecium disparity seems to arise from lower spatial constraints to the production of variable numbers of stamens (Chartier et al., 2017). This additional source of variation, which was not captured here, could have raised the disparity of *Mimosa* androecium in our analyses, as stamens can be equal or twice the number of petals (Barneby, 1991). Nonetheless, this trait is relatively constant among taxa and shows few transitions across the *Mimosa* phylogeny (Barneby, 1991; Simon et al., 2011).

Finally, the innermost floral whorl, the gynoecium showed intermediate level of disparity. *Mimosa* gynoecium, as in the majority of legumes, is monocarpellate (Endress, 2011) and commonly comprised by a compressed ovarium, a filiform style, generally as long as the filaments, ending in a porate stigma (Barneby, 1991; Borges et al., 2024). Even though these traits are generally stable in *Mimosa*, some lineages show styles notably longer or shorter than the filaments (Barneby, 1991), while others undergo carpel abortion during development, resulting in flowers with a rudimentary and non-functional style (Gonçalves et al., 2024). As here we only measured hermaphrodite flowers, the variations in style length may explain the intermediate levels of disparity we found for the gynoecium. Alternatively, it is possible *Mimosa* gynoecium is under the same evolutionary pressures that shape intermediate levels of disparity for other angiosperms (Chartier et al., 2017; Herting et al., 2023).

Taken together, these patterns suggest that development associates with the disparity of *Mimosa* floral whorls. The greater developmental variability of the calyx is linked to higher levels of disparity, while the limited morphological variation of the three other whorls reflects their more canalized development.

The limitation to floral evolution is reinforced by all whorls and their parts evolving around a phenotypic optimum (Table 1, Table S3), as former indicated by the lack of clear and differentiated regimes structuring phylomorphospaces (Figure 2; e.g., constrasting with Dellinger et al., 2019; Kriebel et al., 2022; Reich et al., 2020). The gradient of half-life and stationary variance values across floral whorls seen here adds weight to the idea that they are subjected to different levels of constraint, as highlighted by the development patterns discussed above. Evolutionary constraints over traits of the reproductive whorls (i.e., androecium and gynoecium) are particularly more intense, as evidenced by lower values of half-life and stationary variance, suggesting a strong and fast pull towards an optimum morphology. In other groups, lower half-life values were associated with rapid transition between pollination systems (Joly et al., 2018), which, as discussed above, does not seem to be the case for the less variable *Mimosa* flowers. Moreover, linking the phenotypic diversity of these flowers to that of pollinators is problematic because these animals are mostly attracted to and interact with the inflorescence as a whole. If *Mimosa* inflorescence acts as the main unit of reproduction, then it should also play a role in the evolution of individual flowers.

### Inflorescence functioning canalizes the evolution of floral traits

When flowers are densely arranged in inflorescences, developmental and functional constraints exerted by this hyperstructure may canalize the evolution of individual floral traits. For example, changes in floral symmetry, one of the most remarkable aspects of floral diversity, may be induced by interplay between neighboring floral meristems within densely packed inflorescences. (Baczyński et al., 2022; Naghiloo, 2020; Ronse De Craene, 2018). In the Asteraceae capitulum, and in the peculiar inflorescence of *Parkia*, another mimosoid legume, distinct floral morphotypes arise from temporal shifts in development induced by changes in the spatial arrangement of flower clusters (Dadpour et al., 2011; Moraes et al., 2025). Although *Mimosa* flowers are mostly homomorphic (Barneby, 1991), spatial constraints related to the organization of the inflorescence may also affect the diversity of floral traits. For example, variation in petal number is more frequent in congested inflorescences (Gonçalves et al., 2024) and also seen in *Mimosa* (Barneby, 1991; Borges et al., 2024). These traits are more free to evolve without negative impact on reproductive success because they do not directly alter the structure of the *Mimosa* inflorescence, the actual unit of attraction and interaction with pollinators. On the other hand, changes in perianth length require a corresponding change in the length of reproductive organs to maintain the open morphology of the flower and, thus, form and function of the brush-like mimosoid inflorescence. Accordingly, we showed that, across all whorls, traits describing vertical variation are less disparate and more strongly maintained around a morphological optimum than traits describing horizontal variation. This pattern is evident in values of both half-life and stationary variance of individual floral traits (Figure 5). Taken together, these results suggest that floral traits that directly shape the inflorescence, such as filament and style length, are the most evolutionarily constrained. On this ground, we suggest that the patterns of floral uniformity and convergence highlighted here are underlaid by (developmental) constraints imposed by the form and function of the inflorescence in *Mimosa*.

## Conclusion

Despite being key components of the world’s flora, our knowledge of the macroevolution of inflorescences and flowers with uniform morphology still lags behind that of morphologically and ecologically specialized groups. As a result, data, such as those on pollinator systems, are scarce, affecting direct tests of how pollinators may influence morphology in groups with an uniform floral bauplan. Nevertheless, morphology provides an interface between external factors (e.g., pollinators) and internal processes (e.g., development), allowing us to discuss the underlying constraints that shaped floral morphological diversity. Here, by modeling the morphological evolution of flowers in the mega-diverse genus *Mimosa*, we showed that floral morphology of different lineages is likely marked by convergence. While stabilizing selection mediated by pollinator homogeneity explains low floral diversity in other groups, the little available data suggest that pollinators play a limited role in the evolution of *Mimosa* flowers. Instead, floral disparity in the genus is better explained by patterns of floral development within each whorl. Because *Mimosa* flowers are displayed and function within the inflorescence, their morphology must be coordinated with the general configuration of the inflorescence. We therefore suggest that form and function of the inflorescence canalize floral development, and thus the disparity of floral traits contributing the most to inflorescence organization. As macroevolutionary investigation of groups with uniform floral morphology expands, we may unveil new insights into the patterns and processes that have shaped the diversity and evolution of key components of the world’s flora. Here we suggest that low variation in the flowers of these plants may come from convergences mainly driven not by adaptive evolution, such as stabilizing selection, but by (developmental) constraints to morphological change.

## ACKNOWLEDGMENTS

We thank Facundo Martín Labarque for providing access to the microscope and photographic equipment used for collecting morphological data (FAPESP, processes numbers 2017/17263, 2018/22762-3 and 2015/22000-8). We thank Rafael Fernandes Barduzzi for support and comments that improved the quality of this manuscript. This study was financed in part by the Coordenação de Aperfeiçoamento de Pessoal de Nível Superior - Brasil (CAPES) – Finance Code 001, by the São Paulo Research Foundation (FAPESP), Brasil (process numbers 2021/12607-3, 2024/00265-9, 2022/03046-0, 2021/13031-8 and 2023/16170-4) and New York Botanical Garden (Rupert Barneby Award 2022).

## AUTHOR CONTRIBUTIONS

MM, FAM and LMB conceptualized and designed the project. MM collected morphological data and conducted the analyses. MM, FAM, YBS, and LMB contributed to data interpretation and writing, reviewing and editing of all versions of this manuscript. MM and LMB obtained financial support. FAM and LMB contributed equally to this work and share senior authorship. All authors approved the last version of the manuscript.

## DATA AVAILABILITY

The data that support the findings of this study are openly available in GitHub at github.com/moniquemaianne/ms-constrained_floral_evolution.

